# Fast-cWDM Brain MRI: Fast Conditional Wavelet Diffusion Model for Synthesis Brain MRI Modality

**DOI:** 10.64898/2026.02.14.705904

**Authors:** Timothy Sereda, Lina Chato

## Abstract

In this paper, we present a novel and efficient framework for cross-modality medical image synthesis, developed for BraSyn–Task 8. Our method combines the fast-sampling capabilities of the Fast-Denoising Diffusion Probabilistic Model (Fast-DDPM) with Discrete Wavelet-Transformed components, as used in Conditional Wavelet Diffusion Models. By reducing the number of denoising steps to 100 and using wavelet-transformed inputs, we accelerate both training and inference and reduce memory usage while preserving high image quality. The framework was trained on the BraTS 2025 dataset, which includes four magnetic resonance imaging (MRI) modalities: T1-weighted, contrast-enhanced T1-weighted (T1c), T2-weighted, and FLAIR. We developed four independent models, each synthesizing one missing modality from the remaining three. Evaluation on the BraSyn 2025 Task 8 public validation set demonstrated competitive performance using standard image metrics: mean squared error, signal-to-noise ratio, and structural similarity index. Our method achieved Third place in the challenge in the final test data, with fast inference times (average 41– 67 seconds per case). To assess clinical relevance, we applied a pretrained nnU-Net segmentation model on the synthesized modalities. Segmentation results yielded high Dice coefficients: 0.877 for the whole tumor, 0.769 for the tumor core, and 0.667 for the enhancing tumor. These results confirm the effectiveness and reliability of our approach for missing-modality synthesis, enabling accurate downstream analysis in high-dimensional medical imaging tasks. Our team in **the challenge is USD-2025-Chato-Sereda (Team ID: 3551654)**.Github link: **https://github.com/tsereda/brats-synthesis**

## 1 Introduction

Medical imaging systems, such as Magnetic Resonance Imaging (MRI), Computed Tomography (CT), mammography, ultrasound, and fundus photography are essential tools for non-invasive medical evaluations. These medical imaging plays a crucial role in diagnosing a wide range of diseases, including brain tumors, stroke, retinal disorders, breast cancer, and others [1,2].

In recent years, numerous research efforts have explored the use of Artificial Intelligence (AI) to improve diagnostic processes and clinical outcomes in healthcare, particularly in the field of medical imaging [3-7].

AI, especially Deep Learning (DL), including generative models, has shown significant potential in enhancing both the quality and quantity of medical imaging data. Applications include image synthesis, denoising, automatic classification, segmentation, and disease detection. These advancements not only assist clinicians in achieving faster and more accurate diagnoses but also enable the development of robust computer-aided diagnostic systems.

One important case use is the identification and segmentation of brain tumors, including primary and secondary tumors. Accurate segmentation of tumor subregions of gliomas typically requires access to multiple MRI modalities, including T1-weighted (T1), T1 post-contrast (T1c), T2-weighted (T2), and Fluid-Attenuated Inversion Recovery (FLAIR) images. The Brain Tumor Segmentation (BraTS) Challenge has become a benchmark for evaluating the performance of automated algorithms in this domain [8]. However, in real-world clinical settings, it is common for one or more of these modalities to be missing due to cost constraints, time limitations, or acquisition errors.

Missing modalities can prevent benefits from the state-of-the-art segmentation models when these models mostly were developed based on multimodal input of either all four MRI modalities, (Tw1, T1wc, Tw2, and FLAIR), or three of them, excluding the T1 modality. To address this challenge, recent research has focused on generating missing modalities using generative models such as Generative Adversarial Networks (GANs) and Diffusion Models [9-11]. These methods aim to synthesize high-quality, clinically realistic images that can substitute missing inputs, thus preserving or even enhancing model performance.

Contributing to the BraTS 2025 Brain MRI Image Synthesis Challenge (BraSyn)-Global Missing Modality Task 8 [13], we propose a **Fast Conditional Wavelet Diffusion Model (Fast-cWDM)** for efficient cross-modality generation of 3D MRI medical images. Our work focuses on developing a generative model that can accurately synthesize missing MRI modalities to support downstream segmentation tasks and improve diagnostic reliability in incomplete imaging scenarios. By integrating the reduced-step sampling of Fast-DDPM [10] with wavelet-based spatial decomposition in cWDM [12], our approach achieves high-quality synthesis with significantly lower memory usage and faster inference, making it well-suited for practical clinical applications.

The remains of this paper are organized as follows: *Section 2* describes the data and explains the proposed synthesis image generated models; *Section 3* presents experiments conducted to develop and evaluate the best model and displays the results; *Section 4* discusses the contributions and sets future directions to enhance the performance.

## 2 Methodology

### 2.1 Multimodal BraSyn 2025 Data for Missing MRI Modality

BraSyn-missing MRI modality task released multimodal MRI training data [13]: BraTS-GLI 2023 dataset [14], and BraTS-METS 2023 [15]. This data consists of 1251 glioma samples, and 238 Metastasis samples. Gliomas are primary brain tumors that arise from glial cells. Gliomas can be both low-grade and high-grade, depending on how aggressive they are. While **brain metastasis** tumors are a **secondary brain tumor** that originate **outside the brain** (in another organ) and spread (**metastasize**) to the brain through the bloodstream or lymphatic system. Each sample in the training data has four MRI modalities, (T1, T1c, T2, and Flair), in addition to a segmentation file. The segmentation file consists of a segmentation mask of the three brain tumor subregions (Peritumoral edema (ED), Enhancing tumor (ET), Necrotic and/or non-enhancing core (NCR/NETC), as well as healthy brain tissues and background), as shown in Figure 1. The size of each MRI modality is (240 × 240 × 155) voxels. To test the developed models, two stages of test are released by BraSyn. In the first test stage, the challenge released validation data that consists of 219 samples of glioma tumors, and 31 samples of Metastasis tumors. The validation data contains all the four MRI modalities, excluding the segmentation file. While the final test stage contains 219 samples of Glioma, 59 samples of Metastasis, and 283 samples of Meningioma [16], excluding the segmentation file and randomly one of the MRI modalities in each sample. Meningioma is a type of primary brain tumor that arises from the meninges. This tumor is a usually benign, slow-growing brain tumor that forms outside the brain tissue but can compress adjacent brain structures.

**Fig. 1.**
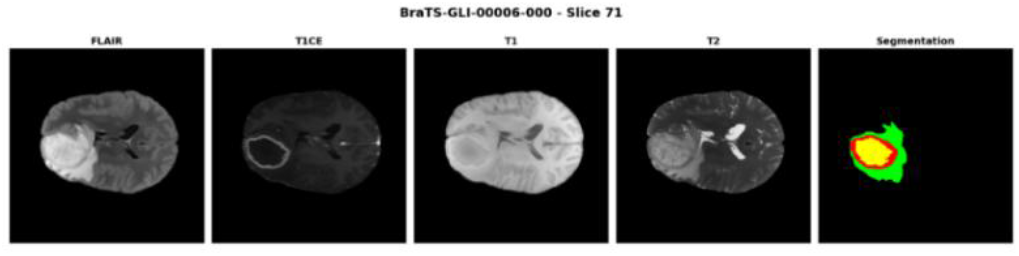
A sample from BraSyn 2025 training data for missing MRI modality task [13]. The colors of the segmentation labels are as follows: Green: ED, Red: ET, Yellow: NCR/NETC.

### 2.2 3D Fast Conditional Wavelet Diffusion Model (3D Fast-cWDM) Background

Denoising Diffusion Probabilistic Model (DDPM) is a type of generative model developed to produce high-quality, realistic images by gradually reversing a noise-adding process [11]. It operates in two main stages:

1. Forward Process: Noise is incrementally added to an image over many steps until it becomes pure noise.
2. Reverse Process: A neural network, such as a U-Net backbone, is trained to denoise the image step by step, reconstructing a clean image from the noisy input.

By learning to reverse this noise process, DDPMs can generate entirely new, realistic images from noise. However, this process usually requires thousands of denoising steps (often 1,000+), resulting in long sampling times. Generating a single image can take several minutes, and the time can increase significantly when generating full 3D/4D medical volumes, depending on computational resources and memory. To address this issue, Fast-DDPM [10] was introduced. The authors proposed a significantly more efficient diffusion process that reduces the number of denoising steps to just 10–25, while preserving image quality. This was achieved through: a) Novel noise schedulers (both uniform and non-uniform), which optimize the diffusion trajectory; b) Aligned training-time sampling, which ensures the model effectively learns denoising within the reduced number of steps. This approach reduced sampling time (by ∼100×) and training costs (by ∼5×), making Fast-DDPM practical for high-dimensional tasks like 3D MRI modality synthesis, without substantial degradation in image quality. Compared to the original DDPM, Fast-DDPM showed only a minor decrease in pixel-wise accuracy (e.g., PSNR, MSE), while maintaining or even improving structural similarity image quality (SSIM), making it well-suited for clinical and research applications. To further accelerate high-resolution medical image generation, the authors of [17] proposed training a diffusion probabilistic model using discrete wavelet components instead of raw images. This strategy offered multiple advantages: 1) Each wavelet component volume is 1/8 the size of the raw image, due to wavelet subsampling reducing spatial dimensions by a factor of 2 in each axis of a 3D image; the reduced input feature map size (1/8 of the original) results in a smaller U-Net, which significantly lowers memory and computational requirements. 2) The U-Net backbone processes these wavelet-based features instead of raw image data, allowing the network to learn from more informative and compact representations. In [17], the authors proposed a **Conditional Wavelet Diffusion Model (cWDM)** for cross-modality synthesis, contributing to the BraTS 2024 missing modality challenge. Their model achieved competitive results compared to standard DDPM. However, the method still involved high computational costs due to the use of 24 wavelet components for the conditional modalities (3 modalities × 8 components for each), in addition to 8 wavelet components for the target (missing) modality. Moreover, since the model was based on standard DDPM, it still required a large number of denoising steps to achieve high-quality outputs. Inspired by Fast-DDPM’s reduced time steps and the efficiency of the cWDM approach, we propose a new framework that combines these techniques for cross-modality generation. Our approach aims to leverage the strengths of both methods: significantly faster inference and lower memory usage without compromising image fidelity, thereby enabling practical and scalable solutions for high-dimensional medical imaging tasks.

#### Modeling

Our proposed model, Fast Conditional Wavelet Diffusion Model (Fast-cWDM), operates in wavelet space, as shown in **Figure 2**, to reduce computational complexity and leverages a reduced denoising step strategy to accelerate sampling.

**Fig. 2.**
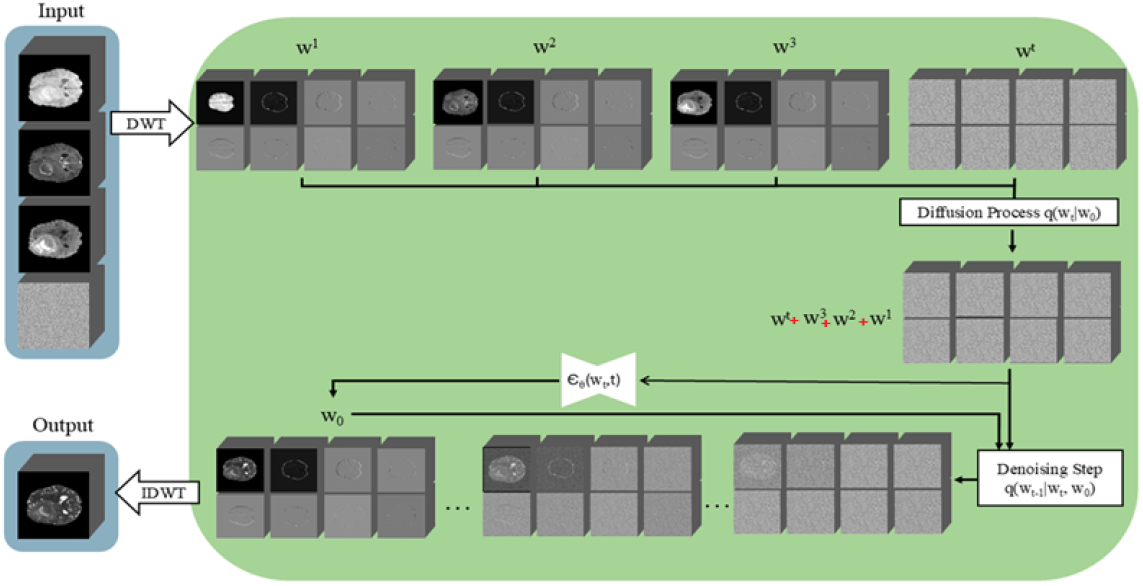
Infrastructure of our Fast-cWDM. The noise image in the input column (left) is the missing modality, and the other available modalities are conditional inputs. The output image is the generated missing image (i.e., noisy image)

We removed 8 pixels from each side of every slice in each MRI modality, and appended 5 additional slices using zero-padding to the final slice of each modality. This preprocessing step ensures a compatible input size of 224×224×160 for wavelet composition. Let the three available MRI modalities *x*^1^, *x*^2^, *x*^3^ ∈ ℝ^224×224×160^, and the target (missing) modality be *x*^*m*^ . Applying the 3D DWT *W*(. ), we obtain

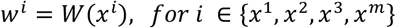

Each *w*^*i*^ ∈ ℝ^112×112×80×*C*^, where *C* refers to the number of the wavelet subbands per modality. As we applied one level of 3D DWT for the volumetric image data, *C* = 8. The model takes {*w*^1^, *w*^2^, *w*^3^} as conditional inputs, and learns to generate *w*^*m*^

##### i. Forward Diffusion Process

We define the forward noising process over the wavelet components of the missing modality. Let *w*_0_ = *ω*^m^, the wavelet representation of the original missing modality. For time step *t* ∈ {1, …, *T*}, the forward process is defined as:

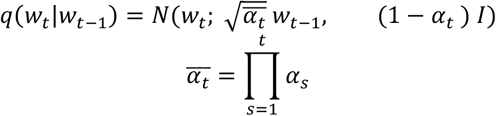

Where N is a Normal Gaussian Noise, *α*_*t*_ = 1 − *β*_*t*_, and *β*_*t*_ is a small noise schedule for our Fast-cWDM (for this task, we used T=100)

##### ii. Reverse Process

The denoising network is a U-Net variant *ϵ*_*θ*_, trained to predict the added noise *ϵ* to the wavelet component of the missing MRI modality, *w*_*t*_, at timestep *t*, and the condition set (the available three modalities) {*w*^1^, *w*^2^, *w*^3^}:

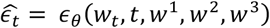

The reverse process uses the predicted noise to reconstruct *w*_*t*−1_ from *w*_*t*_

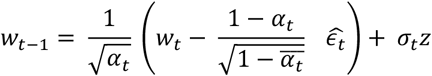

When *σ*_*t*_ is the standard deviation of added noise, and *z* ∼ *N*(0, 1)

The conditional U-Net backbone processes the noisy wavelet component of the missing modality and conditional wavelet inputs. Due to one level of the wavelet subsampling using Haar mother wavelet function, all input volumes are at 1/8 spatial scale, allowing for a lightweight architecture.

### 2.3 Evaluation Measures

To evaluate the performance of our image synthesis model, we adopt a combination of image quality metrics and segmentation-based metrics, reflecting both the visual realism and clinical utility of the generated images.

#### Image Quality Metrics

We assess how closely the synthesized modality resembles the real (acquired) image using the following pixel-level and perceptual metrics:

***Mean Squared Error (MSE)*** quantifies the average squared difference between the synthesized image 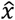 and the ground truth image *x* :

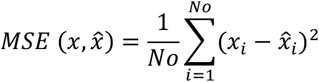

Where No is the number of the image in the test sets. Lower MSE values indicate better pixel-wise accuracy.

***Peak Signal-to-Noise Ratio (PSNR)*** measures the ratio be tween the maximum possible pixel intensity and the distortion (noise) in the image. It is derived from MSE:

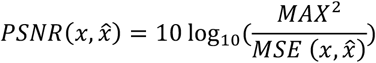

Higher PSNR values indicate better image fidelity.

***Structural Similarity Index Measure (SSIM)*** evaluates perceptual similarity by comparing luminance, contrast, and structure between the synthesized image 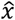 and the ground truth image *x*:

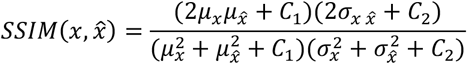

**Segmentation-Based Metrics:** To assess clinical utility, we evaluate whether the synthesized modality enables accurate brain tumor segmentation using a pre-trained BraTS segmentation model. The following metrics are computed between the predicted and ground-truth labels for three tumor subregions:

#### Dice Similarity Coefficient (DSC)

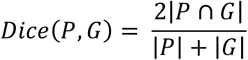

## 3 Experiment setup and Results

We trained a Fast-DDPM model with a 3D U-Net backbone to predict the noise over a reduced number of diffusion steps. We conducted four experiments. For each experiment, we trained a separate model to synthesize one of the four modalities (T1, T1c, T2, or FLAIR) from the remaining three as conditional inputs. Each MRI modality was preprocessed using Haar wavelet decomposition, producing eight subband components per modality, each size 112×112×80. We used T = 100 diffusion steps with a uniform linear noise schedule, trained with the AdamW optimizer (learning rate = 1e-5, batch size = 1) for three stages: 47,500 iterations, 200,000 iterations, and 600,000 iterations using MSE loss. Inference was performed using the Denoising Diffusion Implicit Models (DDIM) solver, exploring uniform timestep sampling strategies. DDIM treats the reverse diffusion process as a deterministic Ordinary Differential Equation (ODE), enabling fast, non-Markovian sampling. All training and evaluation were conducted on four GPUs Nvidia A100 (each 40 GB), that were supported by Nautilus [18]. The U-Net architecture used in all models followed a standard 3D encoder–decoder design with skip connections and group normalization, tailored to handle 8-channel wavelet inputs for each modality. The model output predicted the denoised wavelet coefficients, which were subsequently reconstructed into the full MRI volume using Inverse Discrete Wavelet Transform (IDWT). **Figure 3** displays the training and validation loss for the four missing MRI models. **Table 1** displays the results of image quantitative measures comparing the generated modality against the ground truth (real). All metrics were computed on the reconstructed image volumes in the original spatial domain. nnUnet segmentation model was also used to test the random missing synthesis modality in BraSyn validation dataset, and dice scores of the segmented brain tumor regions are presented in **Table 2**. The qualitative visualization evaluation is presented in **Figure 4**. Some results from the validation data are presented in **Figure 5**.

**Table 1.** Evaluation scores for synthesis MRI images for 250 examples in the BraSyn validation dataset. Number of iterations = 74,500, timesteps =100, validation test conducted 4 times. * Refers to the same models that trained with number of iterations =200,000. ** Refers to the same models that trained with number of iterations =600,000.

| Model | MSE ($\times 10^{-3}$) | PSNR (dB) | SSIM |
| --- | --- | --- | --- |
| **T1** | $2.140 \pm 1.690$ | $27.74 \pm 2.90$ | $0.9386 \pm 0.0234$ |
| **T1c** | $3.189 \pm 2.743$ | $25.93 \pm 2.69$ | $0.9175 \pm 0.0234$ |
| **T2** | $2.665 \pm 1.515$ | $27.35 \pm 3.15$ | $0.9344 \pm 0.0293$ |
| **FLAIR** | $2.532 \pm 3.970$ | $26.58 \pm 2.24$ | $0.9145 \pm 0.0203$ |
| **T1\*** | $1.976 \pm 1.690$ | $28.25 \pm 3.12$ | $0.9399 \pm 0.0224$ |
| **T1c\*** | $2.947 \pm 2.440$ | $26.32 \pm 2.83$ | $0.9159 \pm 0.0246$ |
| **T2\*** | $2.540 \pm 3.834$ | $27.71 \pm 3.32$ | $0.9366 \pm 0.0306$ |
| **FLAIR\*** | $2.336 \pm 1.385$ | $26.90 \pm 2.18$ | $0.9124 \pm 0.0214$ |
| **T1\*\*** | $1.910 \pm 1.630$ | $28.38 \pm 3.09$ | $0.9391 \pm 0.0243$ |
| **T1c\*\*** | $2.994 \pm 2.443$ | $26.23 \pm 2.81$ | $0.9097 \pm 0.0251$ |
| **T2\*\*** | $2.672 \pm 4.231$ | $27.58 \pm 3.37$ | $0.9351 \pm 0.0307$ |
| **FLAIR\*\*** | $2.322 \pm 1.436$ | $26.98 \pm 2.27$ | $0.9126 \pm 0.0226$ |

**Table 2.** nnUnet Segmentation dice scores for the three Brain tumor regions in the BraSyn validation dataset using synthesis generated images for random missing modality. The last row shows the segmentation dice scores for all real MRI modalities.

| Model | WT | ET | TC |
| --- | --- | --- | --- |
| **Fast-cWDM: 100 timesteps, 74,500 iterations** | 0.869 | 0.635 | 0.724 |
| **Fast-cWDM: 100 timesteps, 200,000 iterations** | 0.872 | 0.677 | 0.762 |
| **Fast-cWDM: 100 timesteps, 600,000 iterations** | 0.877 | 0.667 | 0.769 |
| **All MRI modalities are real** | 0.897 | 0.778 | 0.851 |

**Fig. 3.**
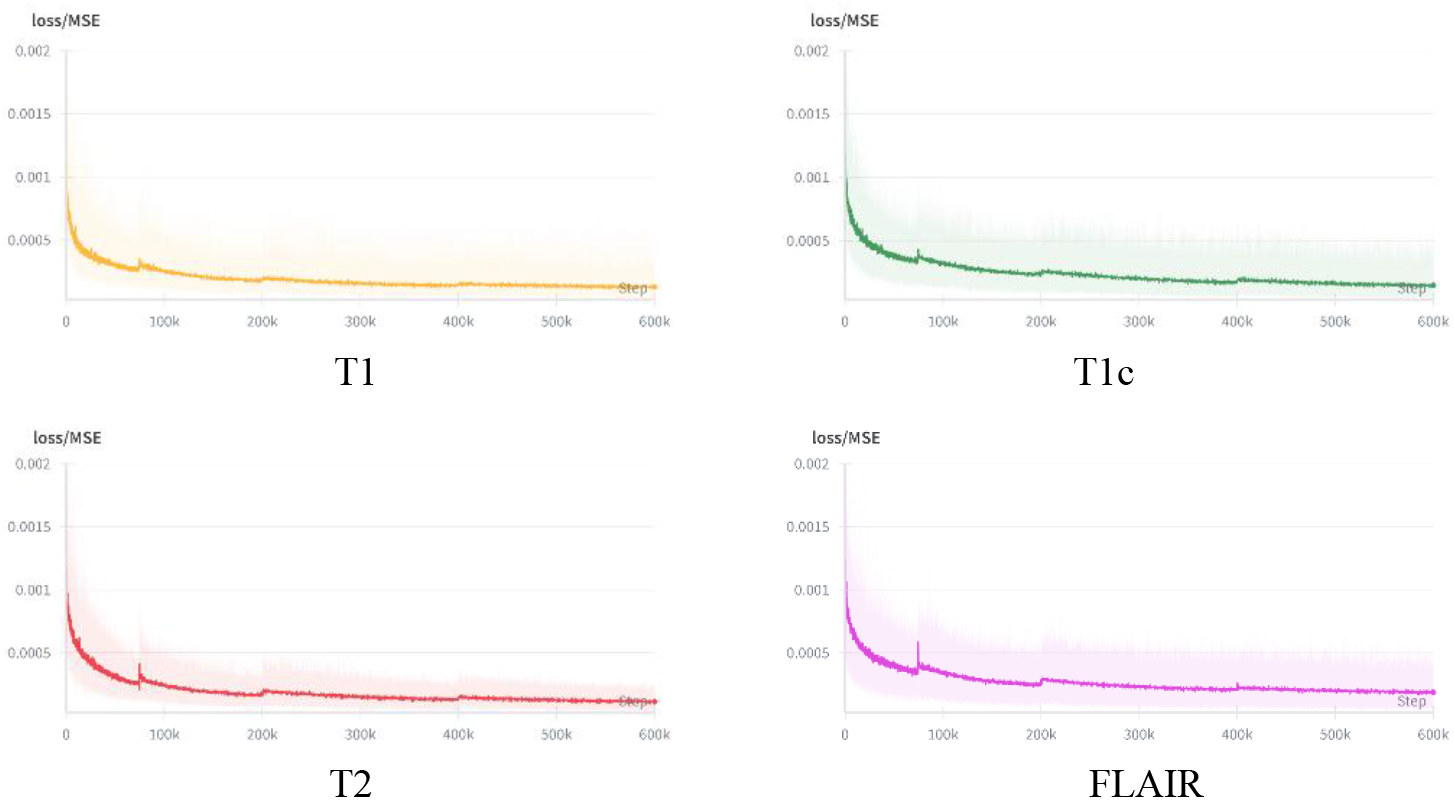
Train/validation loss curves. Each curve is for a specific Fast-cWDM missing MRI modality: T1 (Orange), T1c (Green), T2 (Red), and FLAIR (Purple).

**Fig. 4.**
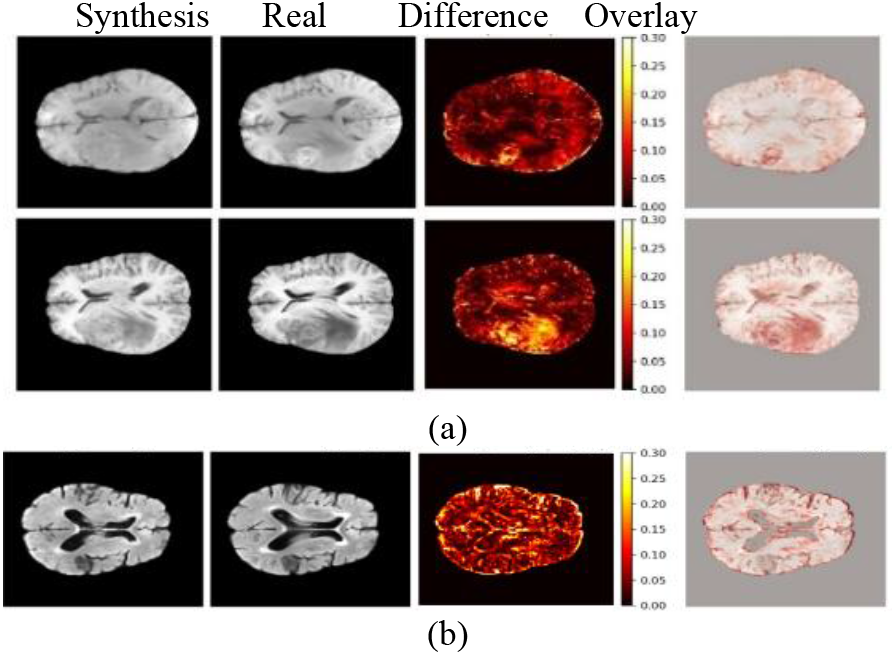
Qualitative comparison of: (a) synthesized T1-weighted MRI and corresponding real image (slice 80) of sample BraTS-GLI-00001-000 (top), and sample BraTS-GLI-00001-001 (bottom); (b) synthesized Flair MRI and corresponding real image (slice 80) of sample BraTS-MET-00207-000.

**Fig. 5.**
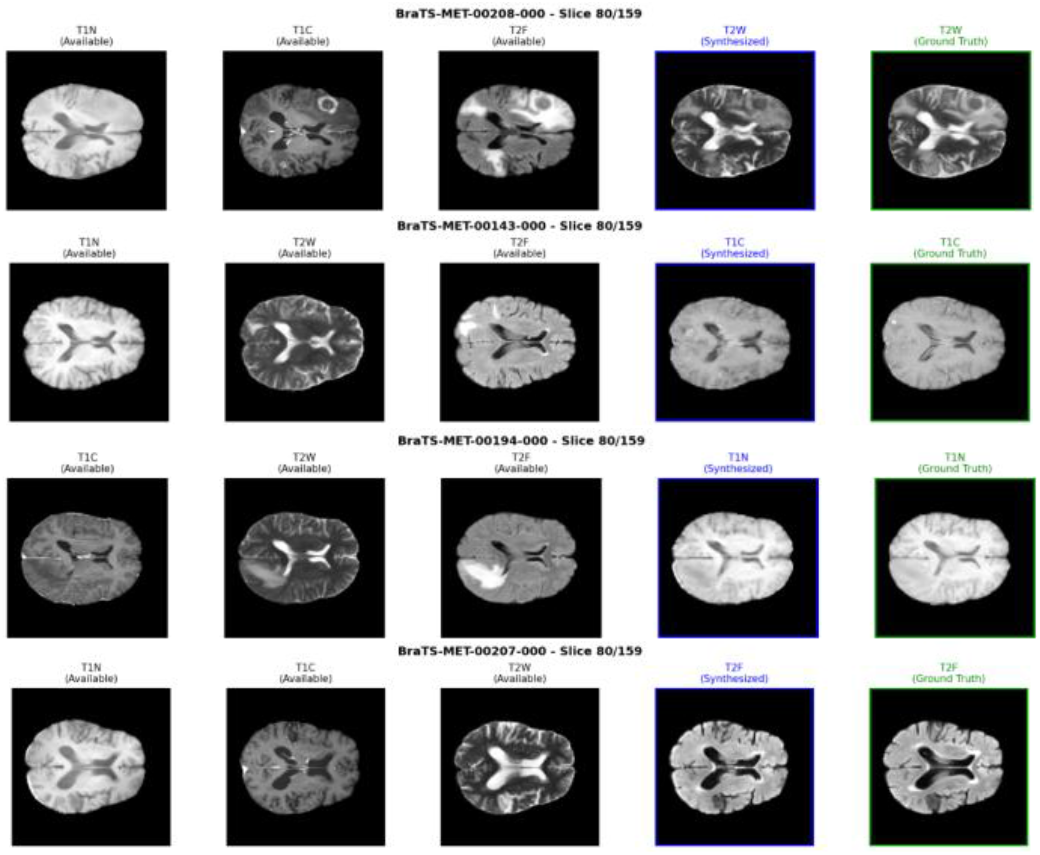
Results from validation dataset.

We submitted the four models (one for each missing MRI modality) that were trained with 600,000 iterations to the BraSyn team 2025 to be tested on the final test phase unlabeled data. **Table 3** shows the training and inference time for the models. **Table 4** presents the results from the final test data using models trained on 600,000 iterations.

**Table 3.**
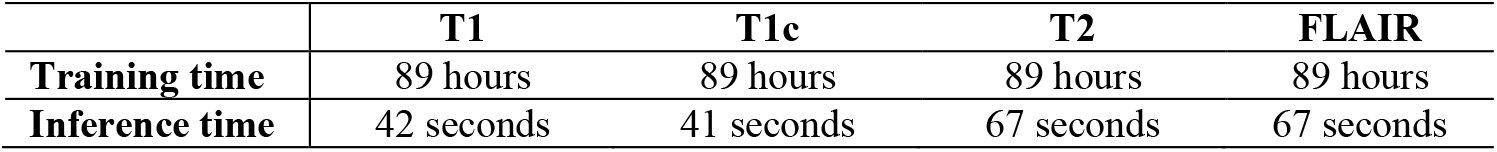
Training time (600,000 iterations) and average inference time/modality for the four Fast-cWDM MRI missing modalities.

**Table 4.** Performance of Fast-cWDM models in the final test data reported by BraTS 2025-Task 8 (achieving 3^rd^ place in the challenge). GLI: Glioma, MEN: Meningioma.

| Group | Type | Dice_ET | Dice_TC | Dice_WT | NSD_0.5<br>ET | NSD_0.5<br>TC | NSD_0.5<br>WT | SSIM |
| --- | --- | --- | --- | --- | --- | --- | --- | --- |
| GLI | Mean | 0.7443 | 0.8073 | 0.9168 | 0.5312 | 0.4685 | 0.5006 | 0.9304 |
| GLI | Std | 0.2866 | 0.2746 | 0.0985 | 0.2887 | 0.2944 | 0.1923 | 0.0510 |
| MEN | Mean | 0.7210 | 0.7349 | 0.7855 | 0.5652 | 0.5638 | 0.5424 | 0.9279 |
| MEN | Std | 0.3836 | 0.3753 | 0.3341 | 0.3513 | 0.3539 | 0.2866 | 0.0223 |
| ALL | Mean | 0.7253 | 0.7730 | 0.8609 | 0.5326 | 0.4931 | 0.5011 | 0.9291 |
| ALL | Std | 0.3251 | 0.3179 | 0.2295 | 0.3094 | 0.3160 | 0.2317 | 0.0427 |

## 4 Conclusion and Future Work

We developed Fast-cWDM, a fast and memory-efficient framework for cross-modality 3D MRI synthesis, contributing to the BraTS 2025 BraSyn Challenge (Task 8) [13-16, 19-21]. By integrating reduced-step Fast-DDPM sampling with wavelet-based decomposition, our method synthesizes missing modalities with high image quality and low computational cost. Our models achieved competitive performance on image quality metrics and yielded strong tumor segmentation results, as shown in **Table 2**. In this stage, we cannot compare our results to the leader board on validation phase as we are not sure if other teams’ segmentation scores were based on nnUnet. These results highlight the practical value of our approach in enabling robust downstream analysis in incomplete imaging scenarios for both clinical uses and AI based diagnosis. From **Table 1**, we can see Fast-cWDM models for T1 and T2 achieved better image evaluation scores compared to other two modalities’ models. Our models, trained for 600,000 iterations, were evaluated by the BraSyn 2025 team on the final test data, and they reported that our submissions achieved **third place** in this year’s challenge. Although the SSIM scores for all groups of the tumors in the final test data, it seems the segmentation dice scores were the best in the glioma brain tumors, as shown in Table 4.

To further enhance our results, we propose training on smaller image patches and experimenting with different timestep schedules, including optimized timestep sampling strategies. In the third column of **Figure 3**, we can see the absolute difference maps between the synthesized T1-weighted MRI slices and the corresponding ground truth images. These heatmaps highlight the spatial distribution of reconstruction errors across the brain slice. Notably, larger differences are observed around structural boundaries and tumor regions, where intensity variations and complex textures increase the modeling challenge. In contrast, homogeneous tissue regions exhibit lower reconstruction errors, reflecting the model’s ability to capture smoother anatomical patterns. This qualitative evaluation helps identify specific areas where synthesis performance could be further improved, such as enhancing boundary precision and tumor representation. However, Flair modality does not show any large differences in the tumor region. Based on the above, we suggest as future work to integrate attention mechanisms to better focus on tumor regions and boundaries, combine synthesis with segmentation using multi-task learning, and use perceptual or adversarial losses to improve detail. Exploring multi-scale architecture or patch-based refinement may also help capture both global and local features for more accurate synthesis. In addition, we plan to explore alternative mother wavelets, such as Daubechies-2 (db2) and Daubechies-4 (db4), which may improve performance over the currently used Haar wavelet. Unlike Haar, which tends to aggressively downsample and emphasize sharp edges, Daubechies wavelets have longer support and smoother basis functions, allowing them to capture more contextual information. This may be particularly beneficial for encoding texture and soft transitions in anatomical structures, potentially enhancing both synthesis quality and downstream segmentation performance.

